# Multimodal genomic markers predict immunotherapy response in the head and neck squamous cell carcinoma

**DOI:** 10.1101/2021.03.30.437691

**Authors:** Gaurav Kumar, Andrew P. South, Joseph M. Curry, Alban Linnenbach, Larry A Harshyne, Adam Ertel, Paolo Fortina, Adam Luginbuhl

## Abstract

Immune checkpoint blockade (ICB) therapy has had a major impact on the clinical management of head and neck squamous cell carcinoma (HNSCC). However, clinical responses to ICB are observed in only a fraction of patients. In the present paper, we used multimodal approaches to a) evaluate the utility of existing ICB biomarkers including tumor mutation burden (TMB), microsatellite instability (MSI), and a T-cell specific signature in predicting therapy response in HNSCC using TCGA cohort, and; b) identify a novel molecular signature to predict ICB therapy response using an ICB clinical trial of HNSCC. Our results confirm previous reports showing TMB efficacy as a biomarker and its outcome can be influenced by age, tumor sub-site, and smoking status; and that High-TMB and high-MSI tumors are associated with T-cell signature, and better survival probability in the HNSCC. We go on to demonstrate that High-TMB and high-MSI utilizes cell-cycle/cell proliferation processes for their molecular functionality; and identify a novel Cell Proliferation ICB-therapy Predicting (CPIP) signature capable of predicting ICB therapy response in HNSCC, retrospectively, where traditional biomarkers of ICB response were insufficient. In summary, the present study defines strategies and novel signatures that can be appropriately used for patient selection for ICB therapy that can improve the clinical outcomes of HNSCC patients.

## INTRODUCTION

Head and neck squamous cell carcinoma (HNSCC) is an aggressive, genetically complex disease characterized by malignant and uncontrolled growth of cells at various sites, including the oral cavity, larynx, oropharynx, hypopharynx, paranasal sinuses, and nasal cavity. Globally, HNSCC is the sixth most common malignancy and accounts for over 650,000 new cases annually with a mortality of approximately 50-60% per year. By far, the commonest site of cancer within this region is the oral cavity, and the commonest type of tumor is squamous cell carcinoma (SCC), which accounts for over 90% of all malignancies at this site [1]. In the United States, HNSCC accounts for 4% of all cancers, with nearly 50,000 Americans diagnosed annually and nearly 11,000 succumbing to their disease [2]. Long-term tobacco use, consumption of alcohol, and infection with high-risk types of Human Papilloma Virus (HPV), are considered the main oncogenic drivers [3].

Immunotherapy, aiming to boost natural defenses to eliminate malignant cells, is a monumental breakthrough for cancer treatment and has revolutionized the field of oncology [4]. It eradicates the tumor cells by inducing or enhancing anti-tumor immune responses in active or passive ways, including adoptive cell transfer (ACT) and immune checkpoint blockades (ICB) [4, 5]. Although immunotherapies demonstrate durable clinical responses, their efficacies vary, and only subsets of patients benefit from them [6, 7]. To augment the clinical benefits of ICB therapies to HNSCC patients, identification of molecular markers/signatures, and approaches predicting therapy response is extremely urgent. These signatures and markers will enable predictions about which patients will respond most successfully to ICB therapies. Identifying those patients who are likely to respond will not only improve their clinical outcomes but also inform about mechanisms of therapy resistance and highlight future therapeutic targets and designs.

Among the emerging biomarkers for ICB response are PD-1/PD-L1 axis, T-cell inflamed signature, tumor mutational burden (TMB), and microsatellite instability (MSI). While their role in predicting ICB response in multiple cancers such as non-small-cell lung cancer (NSCLC), colon cancer, and melanoma [14,18] is well-established, their value in HNSCC is an evolving area of research. At present, majorly, TMB and PD-1/PD-L1 axis are utilized to predict ICB therapy response in HNSCC. However, due to their inconsistency in predicting responses, more robust signatures and approaches are urgently needed [8–12]. On that front, [13, 14] demonstrated success in predicting ICB therapy response in HNSCC using a T-cell inflamed signature. However, as each HNSCC patient can have a variable baseline immune-reactivity due to their variable phenotype and associated risk factors, immune-based predictive molecular signatures might not be sufficient to predict the ICB therapy response. Hence, more robust markers/approaches are needed. In the present study, we identified a novel molecular signature that is rather than fully relying on immune-response, is based on the cell-proliferation processes. Extracted using multimodal approaches (genomic and transcriptomic data), this signature will not only predict the immunotherapy success, but will also lead to elucidation of mechanisms of tumor-intrinsic and immune mechanism in HNSCC. Additionally, we identified/confirmed the role of TMB and MSI in predicting ICB response, along with their association with external factors, as HNSCC is a clinically, histologically, and molecularly distinct cancer. Collectively, the present study will not only congregate the conclusions of existing predictive markers of ICB response, but will also provide a novel signature that can be implemented for ICB therapy success in HNSCC.

## RESULTS

### Tumor mutation burden analysis

As association of TMB with HNSCC using TCGA-HNSC dataset has been studied before [15–17], we first confirmed our results on mutation profile of TCGA-HNSC, and association of TMB with survival prognosis. Here, we found our observations to be in accordance with these studies [15, 16, 18]. Next, we examined the influence of long-term tobacco use, consumption of alcohol, and infection with the HPV16 virus on TMB outcome in the TCGA-HNSC cohort (n=506). We observed that TMB outcome is significantly influenced by the smoking status (smokers: 3.60 mutations/MB vs. non-smokers: 2.14 mutations/MB; p=0.003) but not by HPV status (HPV-positive: 2.80 mutations/MB vs. HPV-negative: 2.43 mutations/MB; p=0.80) as shown in **Fig. 1A.** Since, it is also believed that age, sex, and tumor subsites can also influence the efficacy of ICB therapy response, we next evaluated the influence of these factors on TMB outcome. Here, we observed that older patients (>60 years) tend to have higher TMB rate as compared to young patients (<45 years) (old: 3.85 mutations/Mb vs. young: 1.67 mutations/Mb; p=0.002) **(Fig. 1A)**. Although, associations of TMB with gender and race was observed to be non-significant, tumor subsite was observed to be highly corelated with the TMB outcome. Here, tumors obtained from larynx were observed to have significantly higher TMB as compared to the ones from oral cavity and/or oropharynx as shown previously by our group [19] and further expanded in **Table 1**.

**Figure 1:**
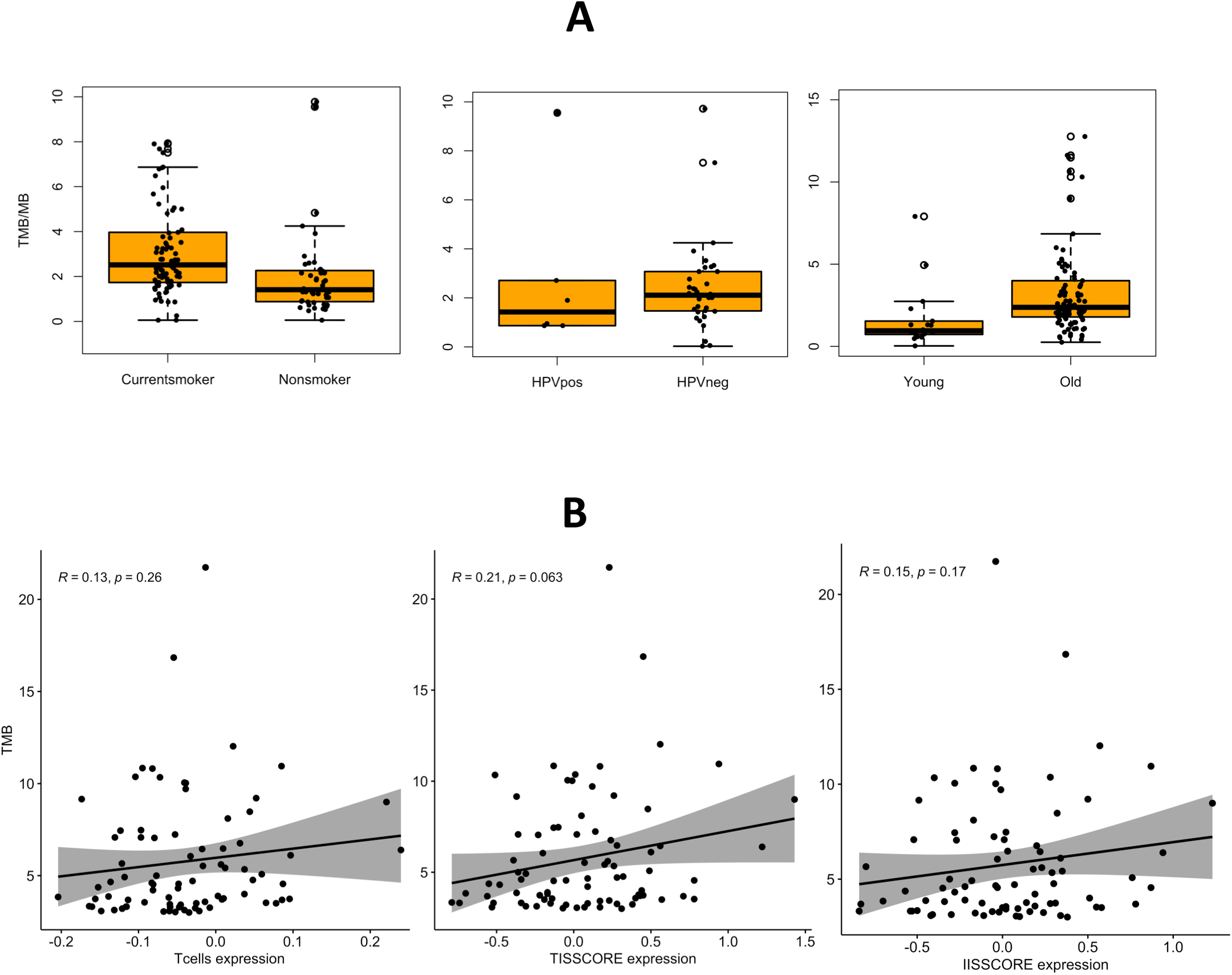
TMB profile in TCGA-HNSC: A) Tobacco smoking status showing significant difference in TMB score between current smokers vs. non-smokers; HPV status showing non-significant difference in TMB status between HPV-positive vs. HPV-negative patients, and; age showing significant difference in TMB score between young (<45 years) vs. old (>60 years); B) Immune-infiltration score (T cells, T-cell infiltration score (TIS), and Immune-cell infiltration score (IIS)) related to TMB is shown to be positively corelated (although non-significant) with higher TMB.

**Table 1:** TMB association with external factors.

|  |  | **TMB**<br>**(mutations/Mb)** | ***P-value*** |
| --- | --- | --- | --- |
| **Sex** | Male | 3.32 |  |
|  | Female | 3.11 | *0.60* |
| **Race** | White | 3.35 |  |
|  | Black or African<br>American (AA) | 2.74 | *0.10* |
| **Site** | Larynx | 4.97 | *larynx vs OC: 0.00001* |
|  | Oral Cavity (OC) | 2.48 | *larynx vs Oropharynx: 0.0002* |
|  | Oropharynx | 2.43 | *OC vs Oropharynx: 0.91* |

It is believed that high TMB tumors contain more neoantigens, that can trigger infiltration of CD8+ T cells stimulating broad antitumor immune responses associated with better clinical outcomes. To evaluate the influence of TMB on regulating the infiltration of immune-specific cells, we observed the correlation of TMB with CD8^+^ T-cells, T cells, and immune cells infiltration. Here, we observed CD8+ T cells, T cells infiltration and immune cells infiltration to be positively correlated with TMB, albeit non-significant **(Fig. 1B)**.

### Microsatellite instability analysis

The MSI score was observed (n=506) to be a mean of 0.19 for TCGA-HNSC cohort. We used this as a cut-off to define high-MSI and low-MSI tumors **(Fig. 2A)**. Here also, first we evaluated the influence of external risk factors on MSI status, and found no significant associations of HPV status, age, sex, and tumor subsite as shown in **Table 2**. Next, correlation of MSI with CD8^+^ T-cells, T cells, and immune cells infiltration was observed. Here, we observed a positive and in fact significant correlation of MSI status with CD8+ T cells, T cells infiltration and immune cells infiltration **(Fig. 2B)**. Finally, impact of MSI on patient’s survival was observed, where, we found patients with higher MSI have better (although non-significant) survival probability as compared to the patients with lower MSI (Chi -square=3.6 on 1 degrees of freedom; log rank p-value=0.06) as shown in **Fig. 2C**.

**Figure 2:**
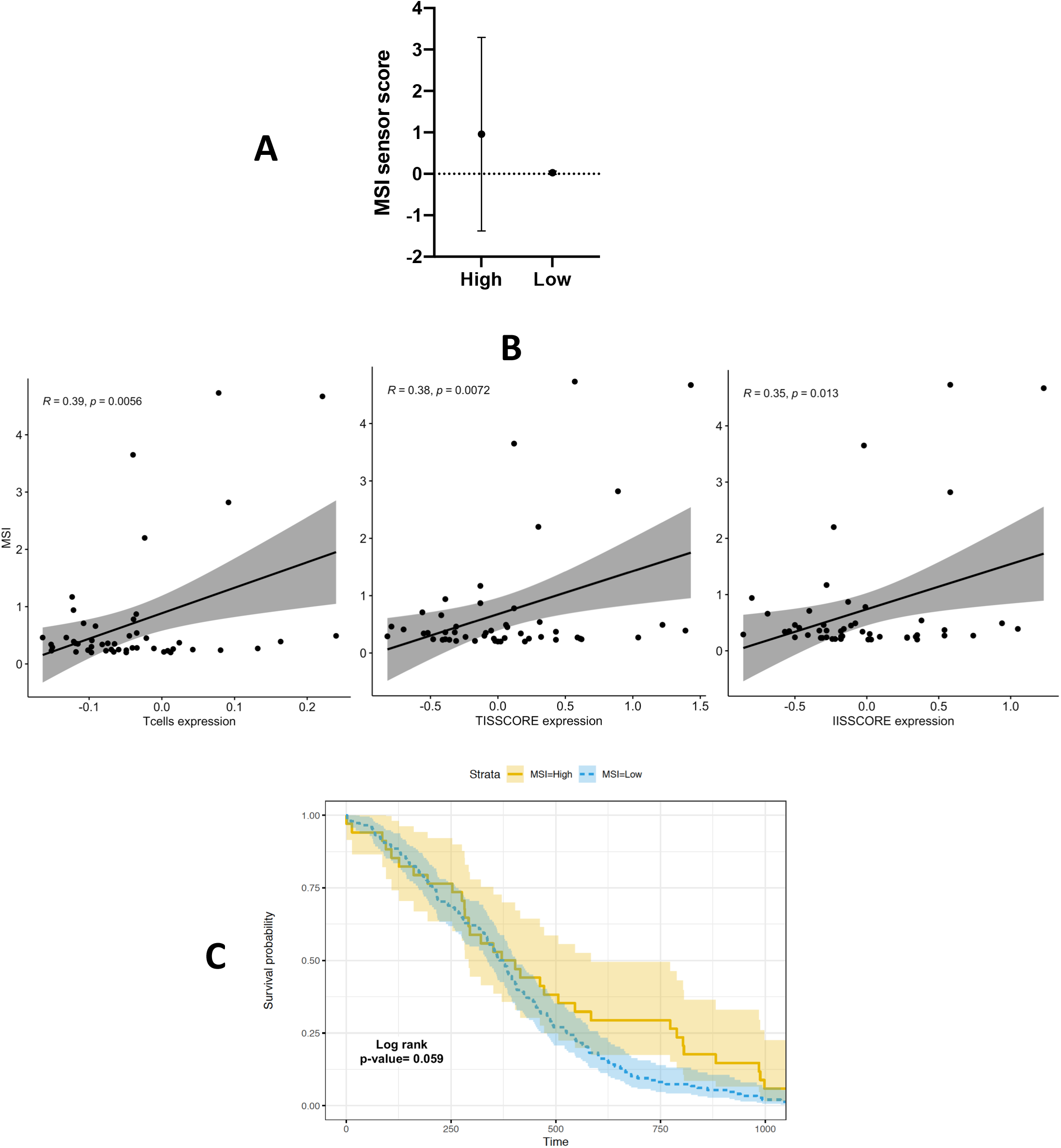
MSI profile in TCGA-HNSC: A) MSI sensor score between high vs low MSI patients; B) Immune-infiltration score (T cells, TIS, and IIS) related to MSI is shown to be positively corelated with higher MSI; C) Kaplan-Meier curve shows better (although non-significant) survival probability of high-MSI HNSCC patients (High-MSI HNSCC patients are represented by the yellow line, and low-MSI patients by the blue line).

**Table 2:** MSI association with external factors.

|  |  | **MSI score** | ***P-value*** |  |  | **MSI score** | ***P-value*** |
| --- | --- | --- | --- | --- | --- | --- | --- |
| **Age** | Young | 0.14 |  | **Sex** | Male | 0.17 |  |
|  | Old | 0.20 | *0.55* |  | Female | 0.13 | *0.64* |
| **Race** | White | 0.16 |  | **Site** | Larynx | 0.14 | *larynx vs OC: 0.94* |
|  | Black or African America (AA) | 0.11 | *0.24* |  | Oral Cavity (OC) | 0.13 | *larynx vs Oropharynx: 0.35* |
| **Smoking** | Life-long non-smokers | 0.25 |  |  | Oropharynx | 0.27 | *OC vs Oropharynx: 0.35* |
|  | Current smokers | 0.18 | *0.64* | **HPV status** | Positive | 0.82 |  |
|  |  |  |  |  | Negative | 0.06 | *0.37* |

### T-cell specific molecular signature

TMB and T-cell-inflamed signature by gene expression profile (GEP) are individually believed to be associated with better prediction of ICB therapy response in multiple cancers, hence, we next tested the correlation of TMB with T-cell specific molecular signature in HNSCC. Here, RNA expression data for high and low TMB TCGA-HNSC cohort was obtained and their ranked gene list was utilized to perform a gene-set enrichment analysis (GSEA) against T-cell specific molecular signature. We observed that high TMB tumors have up-regulated T-cell specific molecular signature in the TCGA-HNSC cohort as shown in **Fig.3A**. This confirms the relationship of TMB and T-cell signature, and their utility in predicting ICB therapy response in HNSCC.

**Figure 3:**
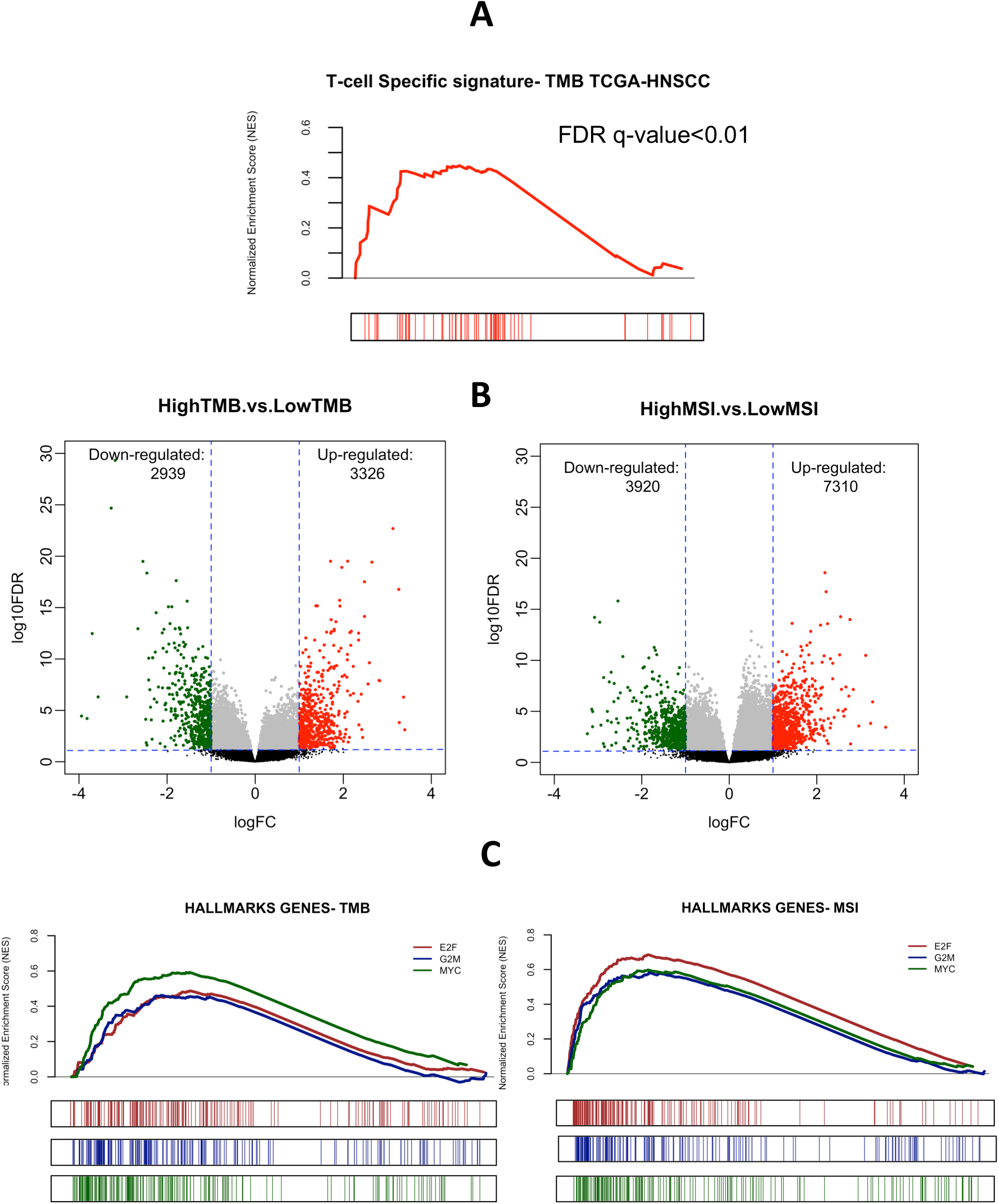
Transcriptomic profile in TCGA-HNSC: A) Gene set enrichment of T-cell specific molecular signature demonstrating high-TMB TCGA-HNSC tumors have high T-cell specific signature; B) Number of differentially expressed genes between high-TMB vs. low-TMB, and high-MSI vs. low-MSI; C) Gene set enrichment of hallmarks of cancer showing up-regulation of cell-cycle and progression processes in high-TMB/MSI tumors.

### TMB and MSI related molecular signature

Next, to identify the molecular mechanism of TMB and MSI status, we analyzed the transcriptomic profile of high vs. low TMB and MSI patients. Here, differential expression analysis between high-TMB (n=134) and low-TMB (n=360) was performed, where, we observed 3326 genes to be up-regulated, and 2939 to be down-regulated in high-TMB patients. Similarly, for MSI status, we observed 7310 genes to be up-regulated (n=86), and 3920 (n=408) to be down-regulated in high-MSI patients as shown in **Fig. 3B.** To evaluate the biological processes being implicated by high TMB and MSI status, we performed GSEA analysis using hallmarks of cancer signatures, where we observed cell-cycle and progression processes including MYC, G2M and E2F to be up-regulated in both high TMB and MSI HNSC tumors as shown in **Fig 3C**.

### Cell Proliferation ICB-therapy Predicting signature

It is known that due to variable levels of immune-response in individuals, T-cell specific signatures alone are insufficient to predict ICB therapy response, hence additional signatures/markers are imperative for better immunotherapy response prediction. Hence, we utilized the differentially expressed genes from high-TMB and high-MSI tumors, along with highly ranked genes from the cell-cycle/proliferation processes, to identify a signature that can be further used to predict ICB response in HNSCC. We obtained a unique list of 31 genes related to cell-cycle/progression processes that can be utilized to predict the ICB therapy response in HNSCC, we termed this list as Cell Proliferation ICB-therapy Predicting (CPIP) molecular signature. To validate the CPIP signature efficacy, we utilized RNA sequencing data of pre-therapeutic tumors from phase-II neoadjuvant trial of nivolumab (PD1-inhibitor) in HNSCC. Originally, this cohort had lower expression of PD1/PDL-1/CD8A/IFNG genes **(Fig 4A)**, and T-cell specific molecular signature in the responder group **(Fig. 4B)**. To retrospectively predict the ICB therapy response in this cohort, we ranked the expressed genes from this cohort to perform GSEA against our CPIP signature. Here, we found that responder patients had high-expression of the CPIP signature prior to the therapy as compared to the non-responders. In this specific cohort, out of a total list of 31 genes, 12 genes (AKR1C3, CDX2, CRLF1, DDX4, FGF19, GRM1, NR5A1, PEG10, PRAME, SOHLH1, TRPM1, and TRPM3) were observed to be driving the ICB therapy response prediction **(Fig. 4C)**. As this signature is derived from genes included in functional processes of TMB and MSI, both these endpoints were observed to be highly corelated with the CPIP signature **(Fig. 5A).** Further, as expected, all genes constituting this signature was observed to be in a strong correlation with each other **(Fig. 5B).** Next, to evaluate the mechanism of how CPIP signature genes are corelating with T-cell activation response, we performed a network analysis using normalized expression of genes of CPIP signature, and CD8a from our clinical ICB cohort. Here, we observed that CD8a is directly or indirectly related with majority of the genes of CPIP signature **(Fig. 6)**. Moreover, we observed that in this cohort CD8a is negatively corelated (blue color edges) with the linked genes of CPIP signature. This demonstrates that CPIP signature can be successfully used to predict the ICB therapy response where patients have lower immune profile.

**Figure 4:**
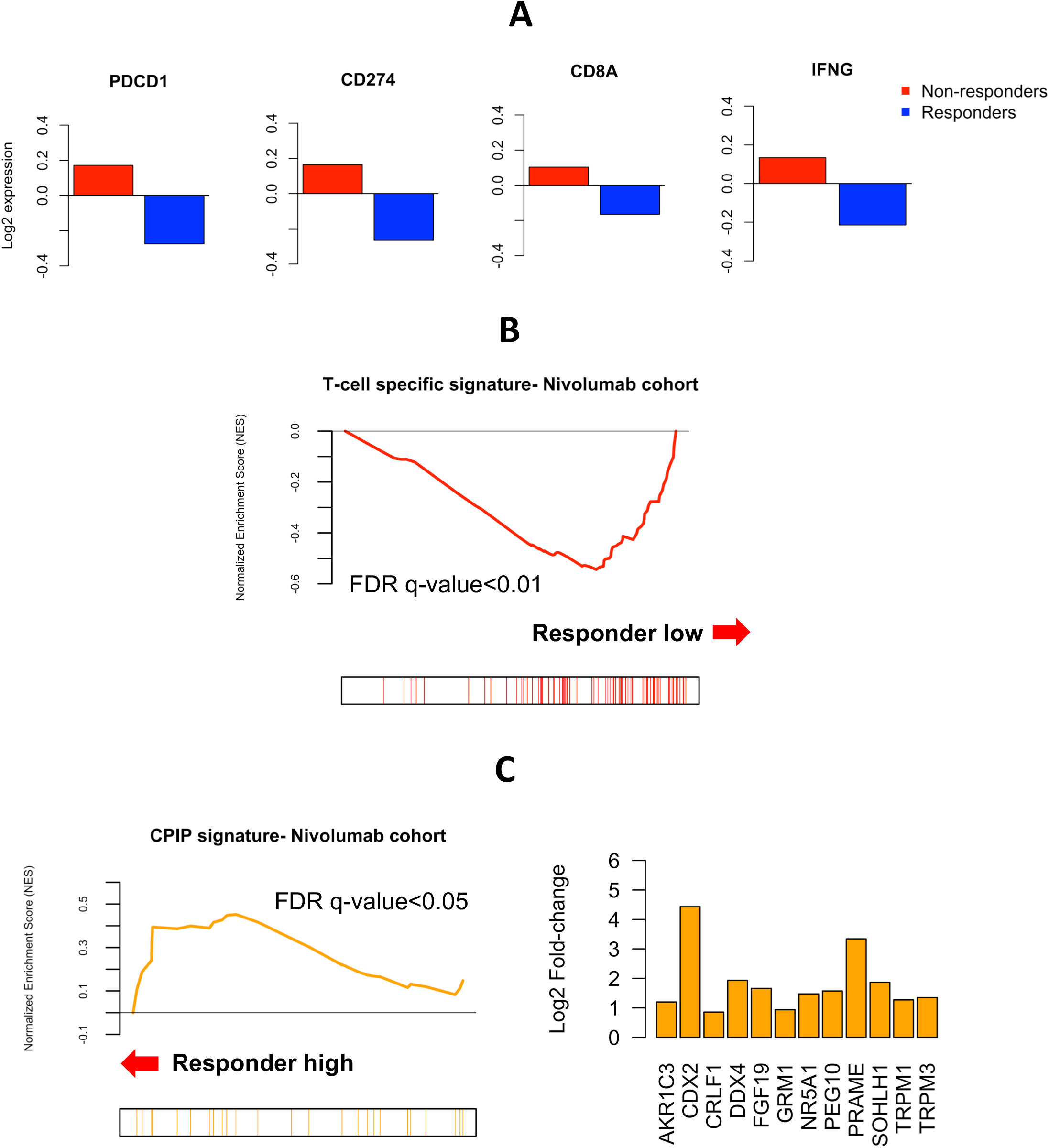
CPIP signature in Nivolumab cohort: A) Reduced log2 expression of established markers of ICB therapy response in the responders from Nivolumab treated HNSCC cohort; B) Gene set enrichment of T-cell specific molecular signature demonstrating responders have low T-cell specific signature; C) Gene set enrichment of CPIP molecular signature demonstrating responders have high CPIP molecular signature, along with log2 fold change of the genes in the leading edge of the analysis.

**Figure 5:**
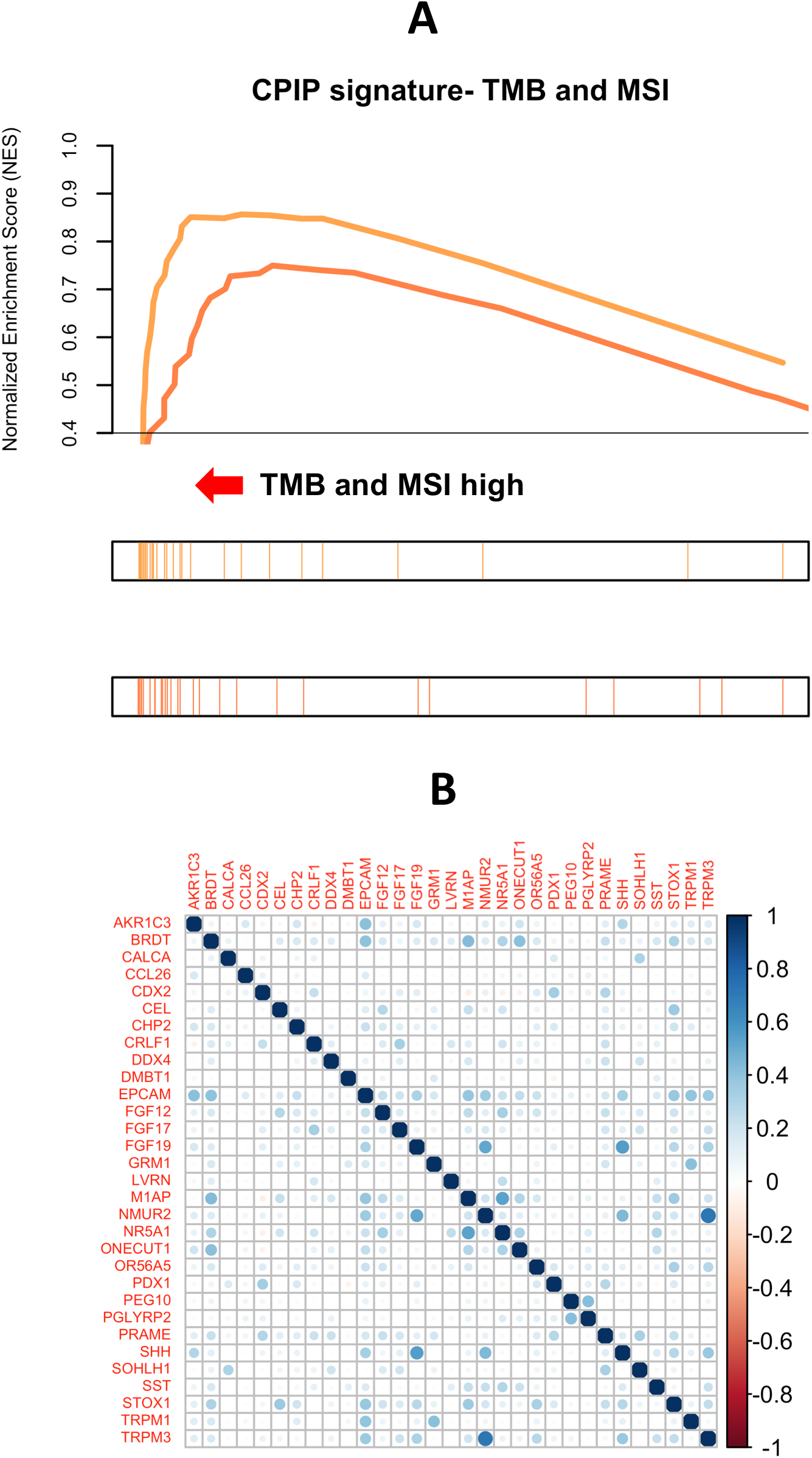
CPIP signature correlations: A) Gene set enrichment of CPIP molecular signature highly corelated with high TMB and MSI tumors in TCGA-HNSC, B) Correlation of all 31 genes of CPIP showing strong correlations between each other.

**Figure 6:**
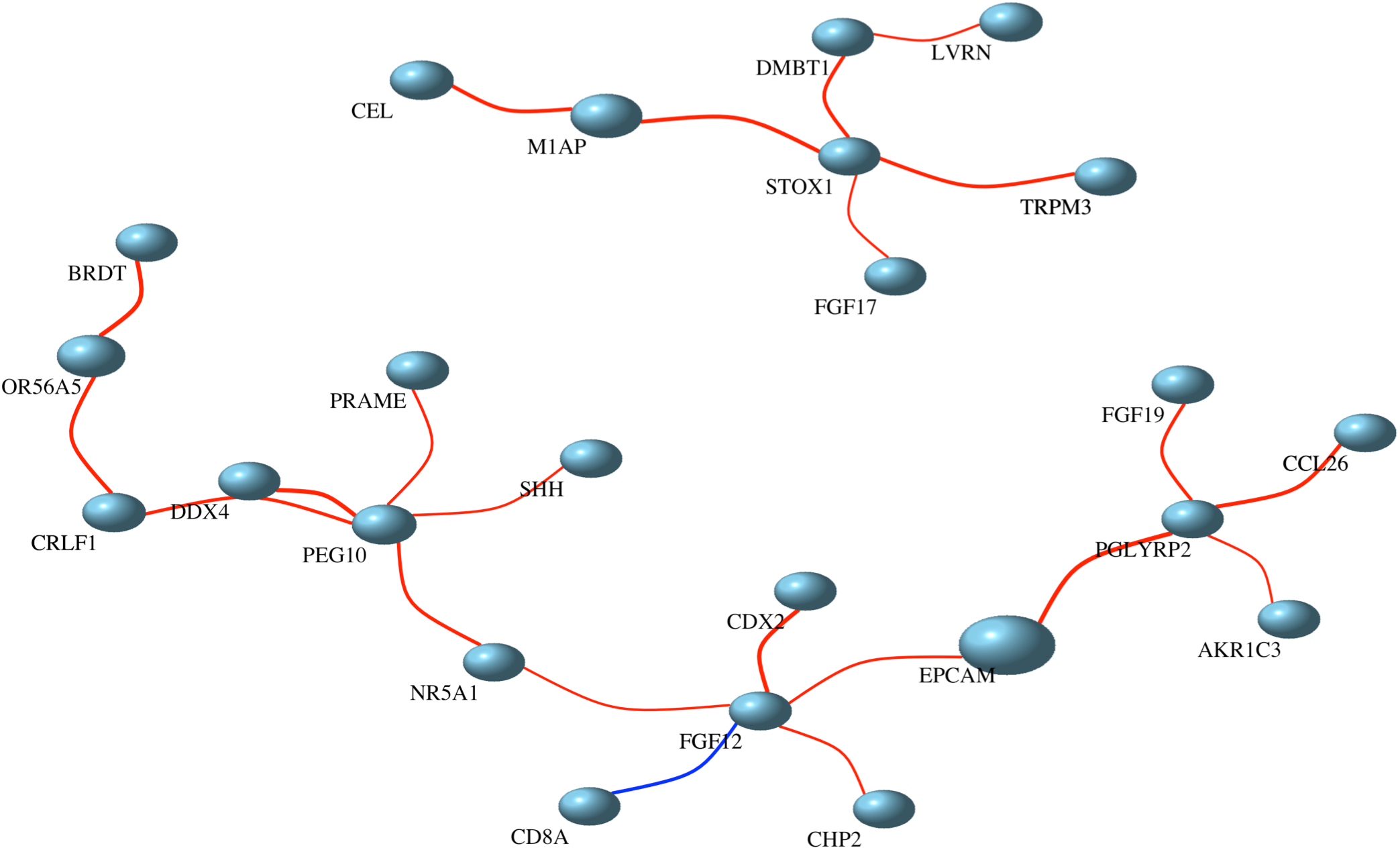
Network analysis of CPIP signature: Network analysis showing interactions of CD8a with the genes in CPIP signature. Here, size refers to their importance in the network due to connection with other genes. Different colors of clusters mean closeness of the related genes. Red color of the edges mean positive correlation and blue color shows negative correlations.

## DISCUSSION

In the present paper, we followed a rationale, step-wise approach to characterize the multimodal genomic markers to predict ICB therapy response in HNSCC.

This study confirms that TMB, and MSI status have strong scientific base as a predictor of immunotherapy response in HNSCC. TMB is a strong genomic biomarker that has been tested in multiple cancers, including HNSCC to predict ICB therapy response. However, since HNSCC is associated with several risk factors, their influence on TMB is not fully studied. Here, we have observed that TMB outcome can be influenced by several external factors known to play role in HNSCC pathogenesis. One of the major factors influencing the TMB is long-term smoking. Tobacco smoke is a complex mixture of chemicals among which at least 60 are carcinogens. Many of these are thought to cause cancer by inducing DNA damage which, if misreplicated, leads to an increased burden of somatic mutations and hence an elevated chance of acquiring “driver” mutations in cancer genes [15]. Tobacco smoking has been previously associated with TMB in NSCLC [20], lung adenocarcinoma [21], along with HNSCC [15, 19]. Our results are in accordance with the previous results, emphasizing the value of utilizing smoking as a confounding factor, when TMB is used as a predictive biomarker in HNSCC. Additionally, TMB is believed to be a site-specific biomarker with important spatial and histologic differences, as we have found strong association of TMB with tumors located in larynx as opposed to the ones located in oral cavity and oropharynx. Our results are in accordance with [19, 22] in HNSCC and NSCLC. As per the demographic correlation, we observed age to have a strong association with the TMB load. Although, mutations accumulate continuously in somatic cells with increasing age, it is unclear whether the age of patients is a determining factor in deriving benefits from immunotherapy. From our results, it is likely that age of the HNSCC patients can influence the TMB outcome. Our results are in accordance with [23–25], where the authors found high correlation of age and TMB in patients of liver, bile duct, colorectal, pancreatic, melanoma, esophageal, HNSCC, and lung cancer.

Recently [18] demonstrated HPV-positive tumors with significantly lower TMB as compared to HPV-negative HNSCC. Although, we observed similar trend in our analysis, but we observed the differences to be non-significant. This may be due to utilizing different cut-off values to define high and low TMB. While, we used the mean TMB/Mb as a cut-off, [18] used only top 25% patients as high-TMB and bottom 25% as low-TMB. Discordance like this provides a strong case to define a universal cut-off value for each cancer type. It should be noted here that generally >10 mutations/Mb is defined as a high-TMB tumor but as suggested by our data and others that each cancer has different mutation rate [15], this cut-off cannot be used for all cancer types to define a high TMB tumor.

MSI status has been mainly utilized to predict ICB therapy response in colorectal cancer. However, after its implementation of predicting PD-1 response in HNSCC [26], more large-scale studies are needed to verify its reliability and stability in clinical applications. In our study, we observed high utility of MSI as a predictor of immunotherapy response. It was observed to have better association with the T-cells activation compared to TMB. This is due to the fact that high TMB/MSI tumors harbor both a higher rate of mutant neo-antigens and a higher number of somatic mutations (including frameshift mutations) that triggers recognition of the tumor by the immune system and result in higher infiltration of T cells leading to improved response to immunotherapies [27]. Interestingly, none of the major external factors were observed to be influenced in our study. Additional molecular studies might be required to further confirm this finding. Using mean cut-off value, MSI survival statistics between high-MSI vs. low-MSI was observed to be on borderline for significance. This might be due to the use of mean as the cut-off. Previous studies [28] have used higher MSI (>4) as a cut-off and have obtained better and significant survival probabilities.

Due to the complex nature of cancer, and imperfection in measurement of the existing predicting biomarkers, new markers and approaches are needed to improve the ICB therapy response in HNSCC. On novel markers front, we identified a molecular signature, CPIP that can successfully predict the ICB therapy response, where existing predicting markers (PD1/PDL-1 axis and T-cell specific signatures) were ineffective. As shown in the results, our Nivolumab treated HNSCC cohort had lower expression of these traditional markers and no existing immune-based marker was clearly predicting the ICB response. Here, our CPIP signature successfully predicted the therapy response, retrospectively. As cell cycle/proliferation process is one of the hallmarks in cancer and is associated with somatic mutations, generation of mutant neo-antigens, and activation of immune-response, it could serve to obtain a prediction on ICB therapy response. Our CPIP signature includes 31 cell cycle/proliferation related genes and based on our network analysis utilizing normalized counts of expression, we observed that at least in our cohort CD8a is negatively corelated with the genes in CPIP signature. Thus, the present study demonstrates that this signature would be very useful to predict the response of the patients that have overall lower immune signature.

On novel approaches front, this study demonstrated that utilizing a gene signature is more robust for therapy prediction instead of using a specific gene. Typically, expression of PD1 and/or PD-L1 axis is used to predict the ICB therapy response, however, they are observed to be imperfect. This inconsistency may be due to several reasons: 1) no specific cut-off value: they are continuous marker with a range of expression levels that complicates its evaluation, 2) technical variation: different assays can generate variable scoring criteria and different thresholds; 3) intratumoral variation: they are a heterogeneous marker, with intratumoral and temporal variation of expression [29]. In the present study, we rather used a molecular signature that is more robust to predict the therapy response. It has already been demonstrated by [13, 14], that a molecular signature is better than a single gene/marker to predict the immunotherapy response.

We note here that further validation of CPIP signature using other published ICB trials, where the immune-signature is not sufficient to predict the therapy response would be ideal. However, access to those studies might be challenging. In future, we plan to expand our cohort and implement machine-learning algorithms to utilize this signature in individual patient settings for ICB therapy prediction in clinical settings.

## CONCLUSION

Our data implicate that TMB, MSI, T-cell specific signatures are strong markers that can be implemented in the HNSCC as a predictive marker for immunotherapy. Our novel CPIP signature can be utilized as a predictive molecular signature for ICB therapy in HNSCC, where immune-signature cannot predict the therapy response.

## METHODS

### TCGA-Data source

The whole-exome sequencing (WES) data in level 3-curated Mutation Annotation Format (MAF) file, RNA sequencing data as raw count matrix, and clinical data for 506 patients of HNSC from The Cancer Genome Atlas (TCGA) was obtained using ‘TCGAbiolinks’ package in R/Bioconductor [30, 31]. The MSISensor data for the TCGA-HNSC was obtained from [28].

### Tumor mutation burden

The TMB load for each cancer type was analyzed by calculating the total number of somatic nonsynonymous mutations per coding area of a tumor exome, described by [32]. The mean TMB was used as the cut-off to determine high vs. low TMB. Mutational analysis was observed using maftools package in R/Bioconductor [33].

### Immune landscape analysis

The immune landscape data including CD8+ T-cells, TIS score, and IIS score was obtained from [6]. Here, TIS was defined as the mean of the standardized values for CD8+ T, T central and effector memory, Th1, Th2, Th17, and Treg cells. IIS was defined as the mean of the standardized values for macrophages, DC subsets, B cells, cytotoxic cells, eosinophils, mast cells, neutrophils, NK cell subsets, and all T cell subsets excluding T gamma delta and T follicular helper cells. T-cell-inflamed gene list was obtained from [14], but was further modified to include additional T-cell-related genes from [6].

### Correlation and survival analysis

Correlations of HPV status, smoking status, age, gender, race, tumor location, CD8+ T cells, TIS, and IIS was observed using ‘ggpubr’ package in R/Bioconductor. Pearson method was utilized for the correlations.

### RNA-sequencing data analysis

The RNA-seq data of pre-therapeutic tumors from phase-II neoadjuvant trial of nivolumab (PD1-inhibitor) was obtained as raw count matrix from Squamous cell carcinoma (SCC) group in Sidney Kimmel Cancer Center, Thomas Jefferson University, PA, USA. Response was characterized as pathologic treatment effect where by each tumor was graded by two pathologists to the degree of treatment effect within the primary tumor [34]. Overall, we had sequencing data from n=13 patients, where, n=5 were responders, and n=8 were non-responders. Before differential expression, batch effects or sample heterogeneity was tested using iSeqQC [35]. Differential gene expression was performed between responder and non-responder using the DESeq2 package [36] in R/Bioconductor. Genes were considered differentially expressed (DE) if they had adjusted p value ≤ 0.05 and absolute fold change ≥ 2. All plots were constructed using R/Bioconductor.

### Functional annotations

The DESeq2 test statistic was used as a ranking metric to perform Gene Set Enrichment Analysis (GSEA) in pre-ranked mode, with genes having zero base mean or “NA” test statistic values filtered out to avoid providing numerous duplicate values. GSEA was performed against custom gene list (T-cell specific genes and CPIP signature), and hallmarks of cancer from MSigDB collections [37]. DAVID [38] was performed on list of DEGs between high vs low TMB and high vs low MSI. Network analysis was performed using igraph [39], where normalized expression counts from Nivolumab HNSCC cohort was used as input. Here, Pearson correlation was used to perform the correlation distances.

## Notes

### Competing Interest Statement

The authors have declared no competing interest.

